# Microbiotyping the sinonasal microbiome

**DOI:** 10.1101/549311

**Authors:** Ahmed Bassiouni, Sathish Paramasivan, Arron Shiffer, Matthew R Dillon, Emily K Cope, Clare Cooksley, Mahnaz Ramezanpour, Sophia Moraitis, Mohammad Javed Ali, Benjamin Bleier, Claudio Callejas, Marjolein E Cornet, Richard G Douglas, Daniel Dutra, Christos Georgalas, Richard J Harvey, Peter H Hwang, Amber U Luong, Rodney J Schlosser, Pongsakorn Tantilipikorn, Marc A Tewfik, Sarah Vreugde, Peter-John Wormald, J Gregory Caporaso, Alkis J Psaltis

## Abstract

This study offers a novel description of the sinonasal microbiome, through an unsupervised machine learning approach combining dimensionality reduction and clustering. We apply our method to the International Sinonasal Microbiome Study (ISMS) dataset of 410 sinus swab samples. We propose three main sinonasal ‘microbiotypes’ or ‘states’: the first is *Corynebacterium*-dominated, the second is *Staphylococcus*-dominated, and the third dominated by the other core genera of the sinonasal microbiome (*Streptococcus*, *Haemophilus*, *Moraxella*, and *Pseudomonas*). The prevalence of the three microbiotypes studied did not differ between healthy and diseased sinuses, but differences in their distribution were evident based on geography. We also describe a potential reciprocal relationship between *Corynebacterium* species and *Staphylococcus aureus*, suggesting that a certain microbial equilibrium between various players is reached in the sinuses. We validate our approach by applying it to a separate 16S rRNA gene sequence dataset of 97 sinus swabs from a different patient cohort. Sinonasal microbiotyping may prove useful in reducing the complexity of describing sinonasal microbiota. It may drive future studies aimed at modeling microbial interactions in the sinuses and in doing so may facilitate the development of a tailored patient-specific approach to the treatment of sinus disease in the future.

## MAIN TEXT

Chronic rhinosinusitis (CRS) is a heterogenous, multi-factorial inflammatory disorder with a complex and incompletely understood aetiopathogenesis.^1^ A potential role of the sinonasal microbiome and its “dysbiosis” in CRS pathophysiology has recently gained increased interest. The nature of the microbial dysbiosis and its role in disease causation and progression however remains unclear, with conflicting findings from the small sinonasal microbiome studies published thus far.

We recently reported the findings of our multi-national, multicenter “International Sinonasal Microbiome Study” or ISMS.^2^ This study, the largest and most diverse of its kind to date, attempted to address many of the limitations of the smaller previous studies, by standardizing collection, processing and analysis of the samples. Furthermore, its large sample size and multinational recruitment, meant that it was more likely to capture geographical and centre-based differences if present. A recent meta-analysis of published sinonasal 16S rRNA sequences revealed that the largest proportion of variance was attributed to differences between studies,^3^ highlighting a role for performing a large multi-centre study that employed a unified methodology.

Contrary to the findings of previous studies, our international cohort showed no significant differences in alpha or beta diversity between the three groups of patients analyzed: healthy control patients without CRS and the two phenotypes of CRS patients, those with polyps (CRSwNP) and those without (CRSsNP). The study however revealed a potential grouping of samples as demonstrated on beta diversity exploratory analysis.^2^ Accordingly, we hypothesized that the bacteriology of the sinuses could be categorized into various clusters of similar compositions. We inquired whether these potential groups would aid in describing the sinonasal microbial composition of patients or associate with clinical features. Similar attempts performed on gut microbiota in healthy individuals were termed *enterotyping*.^4^ The clinical relevance of gut enterotypes remain the topic of research, and sometimes controversy. A previous exploration of clusters of sinus microbiota in patients was performed by Cope et al.^5^ in which the authors reported four compositionally distinct sinonasal microbial community states; the largest group of patients were dominated by a continuum of Staphylococcaceae and Corynebacteriaceae demonstrating a reciprocal relationship.^5^

In this manuscript, we attempt “microbiotyping” to explain interpatient heterogeneity of the bacterial communities in the paranasal sinuses, and are the first to describe “sinonasal microbiotypes” across the first large, multi-centre cohort of individuals with and without CRS. We model our analysis on previous attempts of enterotyping the gut microbiome. We then describe the composition of these microbiotypes, explore potential clinical associations and validate microbiotyping on a separate sinus microbiome dataset.

## RESULTS

### Basic characteristics of the study cohort and beta diversity plots

The main ISMS study cohort was described in our previous publication.^2^ In brief, 410 samples were included in the analysis collected from 13 centres representing 5 continents. These samples are distributed along three diagnosis groups as follows: 99 CRSsNP patients, 172 CRSwNP patients, and 139 (non-CRS) controls. Beta diversity ordination plots (of weighted UniFrac and Jensen-Shannon distances) are shown in Figure 1. The plots do not reveal any distinct grouping by disease state or by centre, but on visual inspection show a triangular arrangement suggesting that samples lie on a continuum between three distinct clusters, providing motivation for further analysis.

**Figure 1:**
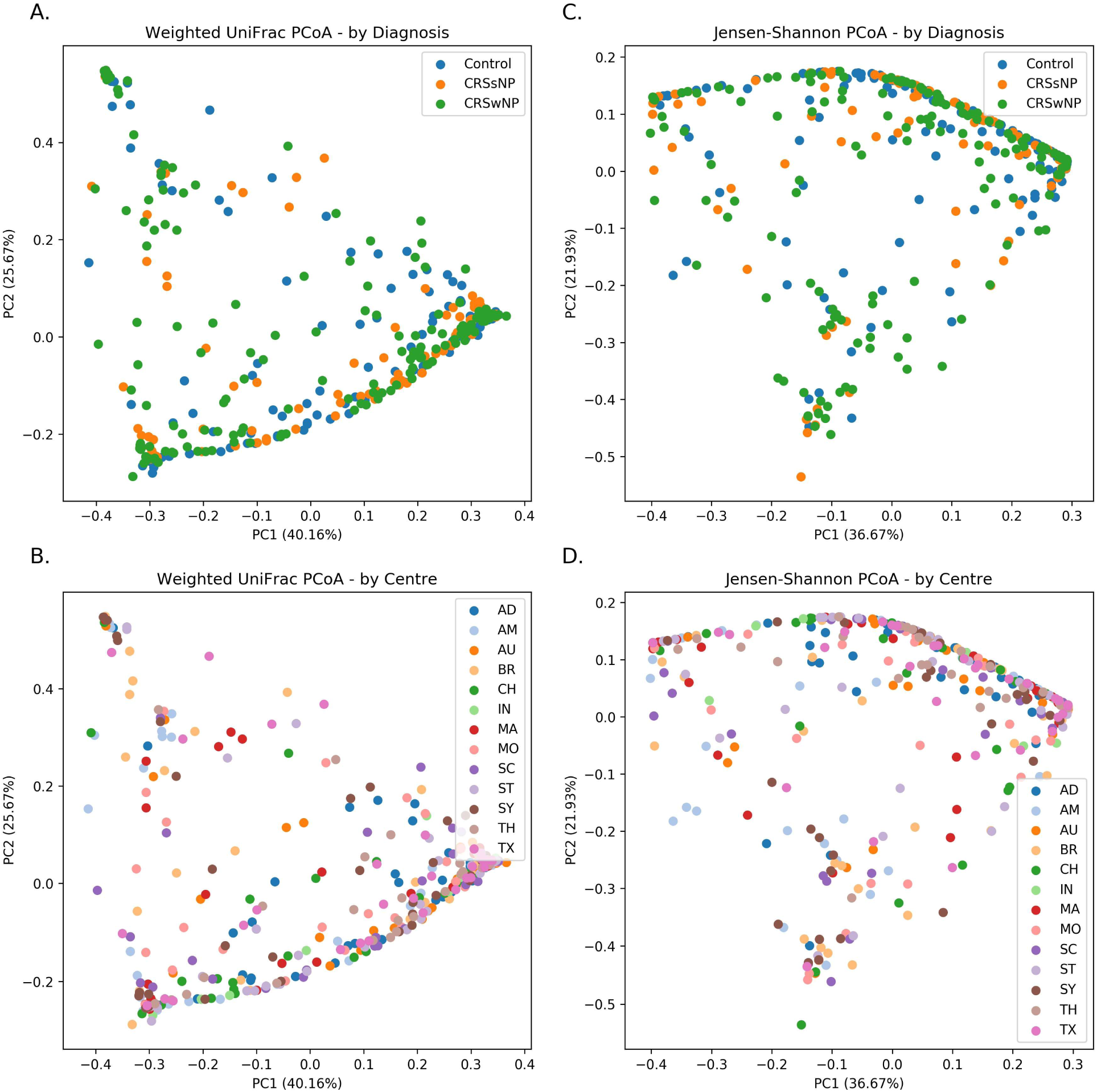
Beta diversity ordination plots.

### Composition of the three sinonasal microbiotypes

We applied our microbiotyping approach through the unsupervised dimensionality reduction and clustering method described in the Methods. The composition of the resulting “sinonasal microbiotypes” is found in Figure 2A.

**Figure 2:**
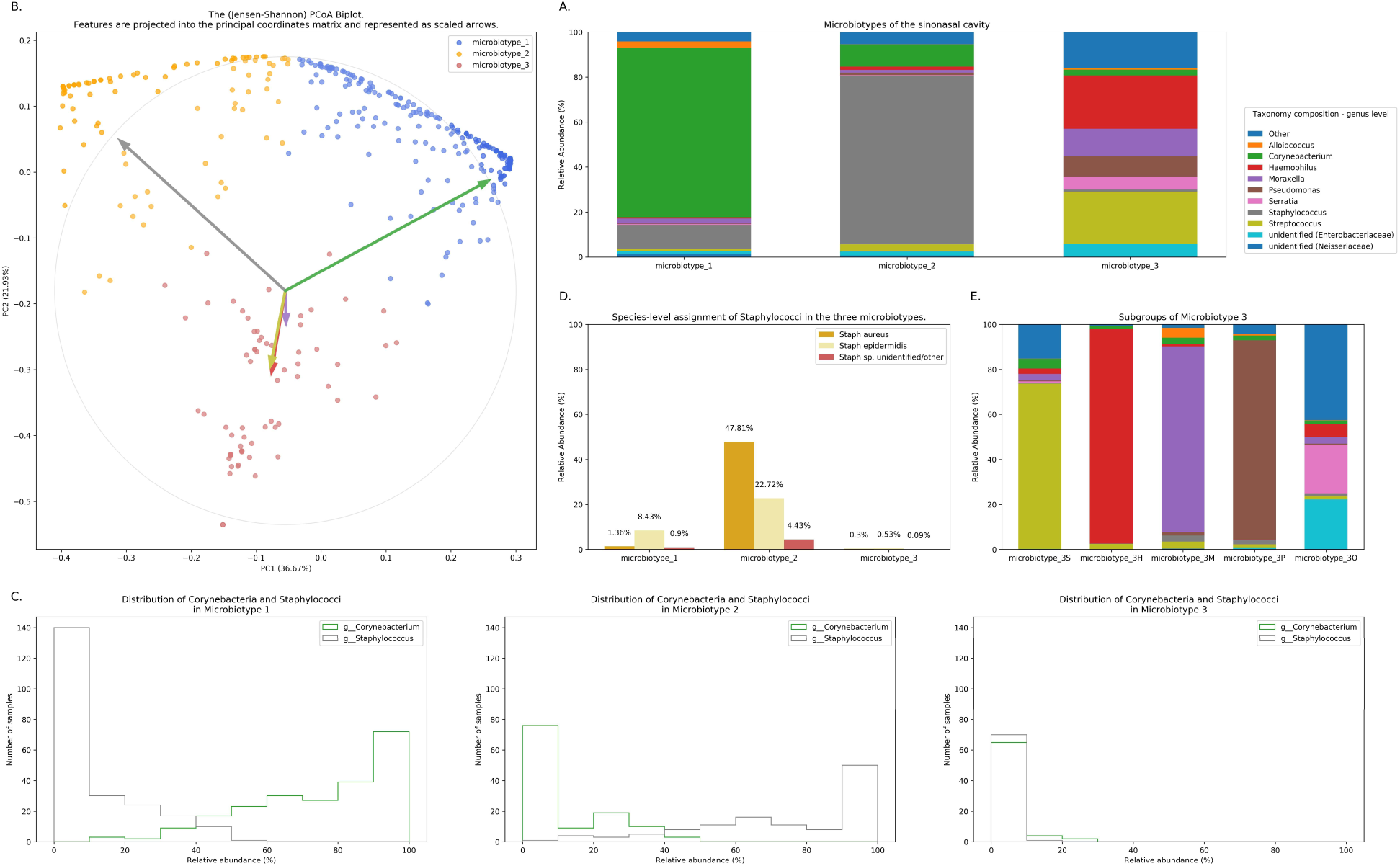
Microbiotyping the sinonasal microbiome. (A) Taxonomic composition of the three microbiotypes at the genus level. (B) Illustration of the assigned microbiotypes on the Jensen-Shannon PCoA biplot. Arrows were used to depict the projection of the genera onto the PCoA matrix. Each arrow is indicated by the color of the genus according to the Legend. (C) Histograms demonstrating the relative abundance of Corynebacterium and Staphylococcus. (D) Distribution of staphylococcal species (mean relative abundance). (E) Subgroups of microbiotype 3 (hierarchical density-based clustering).

Microbiotype 1 is dominated by *Corynebacterium* (mean relative abundance of 75.29%). Microbiotype 2 is dominated by *Staphylococcus* (mean relative abundance of 74.96%). Microbiotype 3 contained samples that were mostly constituted of *Streptococcus*, *Haemophilus*, *Moraxella*, *Pseudomonas* and other genera.

The Abundance/Prevalence tables for the microbiotypes is demonstrated in Supplementary Tables S1A, S1B and S1C.

We used a PCoA biplot to project features (genera) onto the PCoA matrix.^6^ The 5 topmost abundant genera were overlaid on the PCoA plot as arrows, originating from the centre of the plot and pointing to the direction of the projected feature coordinates. (Figure 2B) Each arrow is indicated by the color of the genus according to the Legend in Figure 2A, and the length of each was normalized as a percentage of the longest arrow. The coloring of the samples in 2B in the PCoA scatter plot according to the microbiotype assignment is provided for additional illustration. (Figure 2B) We note that the biplot arrows show a quasi-orthogonal arrangement between the key genera that constitute the microbiome.

The distributions of the relative adundances of *Corynebacterium* and *Staphylococcus* in all three microbiotypes were plotted in histograms (Figure 2C). It was noted that in microbiotype 1, most samples have a high abundance of Corynebacteria (i.e. Corynebacteria dominate), while Staphylococci appeared to dominate in microbiotype 2 in most samples.

### Dissection of “sinonasal microbiotype 3”

We observed that Microbiotype 3 included various genera that did not cluster into the major two microbiotypes. It was also evident that this microbiotype is more heterogeneous. Applying the K-Means algorithm we showed poor clustering on only the first two and three Principal Components, since this group included multiple signatures with various dominant organisms. Accordingly, we employed the hieararchical density-based clustering algorithm “hdbscan”^7^ on the full-dimensional OTU table. One advantage of this algorithm is that it can estimate the number of clusters, without a priori specification by the user. This algorithm also has the ability to detect “outliers” that fail to cluster with the rest of the groups and detaches them into a separate “Miscellaneous/Other” group. We ran this algorithm on samples in Microbiotype 3 and this revealed four clusters, each dominated by one of the genera of *Streptococcus* (21 samples), *Haemophilus* (16 samples), *Moraxella* (9 samples), and *Pseudomonas* (7 samples), with a mean relative abundance ranging from 73.49% to 95.5%. The fifth cluster was the assigned “Miscellaneous/Other” group (18 samples). We term these “sub-microbiotypes”: microbiotype 3S, 3H, 3M, 3P, and 3O, respectively. (Figure 2E)

### Exploring microbiotypes at the species-level reveals potential antagonism between *Corynebacterium* species and *Staphylococcus aureus*

At present, species level assignment is limited by the current technology of 16S-surveys, the current state of microbial databases in general, and by our chosen short-read sequencing methodology. However, species level associations hold clinical significance for sinus health, since *Staphylococcus aureus* has been traditionally associated with biofilm formation and superantigen elaboration, both of which are associated with more severe sinus disease and poorer response to treatment. Furthermore nasal carriage of methicillin-resistant *Staphylococcus aureus* (MRSA) is a global health concern with implications that extend far beyond the sinuses. Moreover, our new QIIME 2-based pipeline^8^ allows a higher “sub-OTU” resolution compared to older pipelines, offering an opportunity to resolve some taxa at species level when possible.^9,10^

We explored taxonomy assignment at the species level, with a focus on Staphylococcal species. Staphylococci were assigned to either *Staphyloccocus aureus*, *Staphylococcus epidermidis* or unclassified *Staphylococcus*. We found that almost all of the assigned *Staphylococcus aureus* species were clustered in Microbiotype 2, forming 47.81% mean relative abundance of this Microbiotype, compared to 1.36% and 0.3% in Microbiotype 1 and Microbiotype 3 respectively. (Figure 2E) Differential abundance of both *Staphylococcus aureus* and epidermidis between the disease groups was confirmed as statistically significant using ANCOM.

In light of this finding, we hypothesized a reciprocal or antagonistic relationship between *Corynebacterium* sp. and *Staphylococcus aureus* and investigated this using SparCC. This confirmed a significant negative correlation between *Corynebacterium* genus and the species *Staphylococcus aureus* (SparCC correlation coefficient = −0.339, p = 0.001). Interestingly, *Staphylococcus* epidermidis positively correlated with *Corynebacterium* (SparCC correlation coefficient = 0.271, p = 0.001). These results should be interpreted cautiously in light of 16S-sequencing limitations. Nevertheless, they do appear to correlate to previous findings in the literature, including in vitro experiments^11^, a murine nasal bacterial interaction model^12^, and a survey of nasal vestibule swabs in healthy individuals^13^. These results suggest that a benign or probiotic role is played by both *Corynebacterium spp*. and *Staphyloccocus epidermidis* when interacting with *Staphylococcus aureus*.

### Prevalence and distribution of the microbiotypes in different diagnoses and centres

Microbiotype 1 was assigned to 222 samples (54.1%), microbiotype 2 to 117 samples (28.5%), and microbiotype 3 to 71 samples (17.3%). The prevalence distribution of the sinonasal microbiotypes did not appear to significantly differ by the disease state of the sinuses. (Figure 3) However, a Chi-Squared test on the contingency table by centre showed significantly different distributions by centre (FDR-corrected p < 0.001): there was a higher prevalence of microbiotype 2 in our European centre (Amsterdam), and a higher prevalence of microbiotype 1 in Asian and Australasian centres, with a much lower prevalence of microbiotype 3 in Asia. (Figure 3 and Table 1)

**Figure 3:**
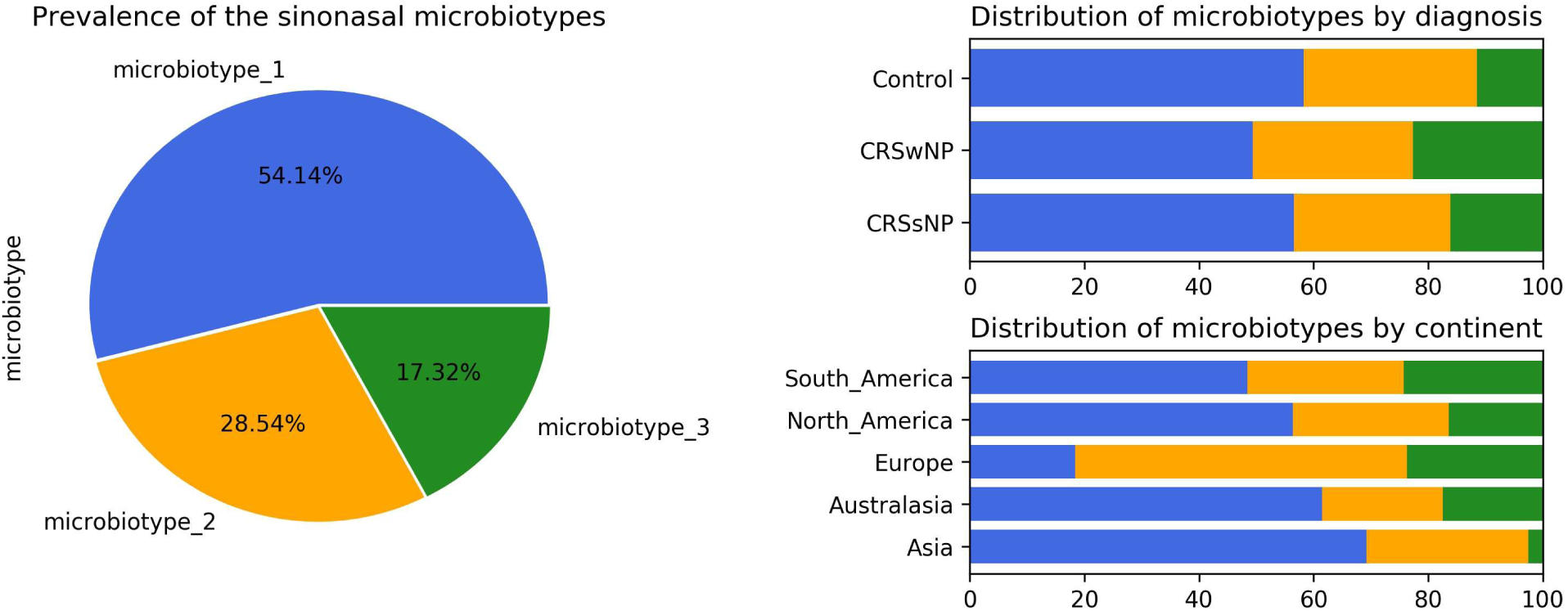
Prevalence and distribution of the microbiotypes.

**Table 1:** Distribution of microbiotypes by diagnosis and continent.

| variable | value | microbiotype_1 | microbiotype_2 | microbiotype_3 | p value |
| --- | --- | --- | --- | --- | --- |
| Diagnosis | CRSsNP | 56 (56.6%) | 27 (27.3%) | 16 (16.2%) | 0.507 |
|  | CRSwNP | 85 (49.4%) | 48 (27.9%) | 39 (22.7%) |  |
|  | Control | 81 (58.3%) | 42 (30.2%) | 16 (11.5%) |  |
| Continent | Asia | 27 (69.2%) | 11 (28.2%) | 1 (2.6%) | < 0.001 |
|  | Australasia | 67 (61.5%) | 23 (21.1%) | 19 (17.4%) |  |
|  | Europe | 7 (18.4%) | 22 (57.9%) | 9 (23.7%) |  |
|  | North_America | 89 (56.3%) | 43 (27.2%) | 26 (16.5%) |  |
|  | South_America | 32 (48.5%) | 18 (27.3%) | 16 (24.2%) |  |

### Associations of microbiotypes with clinical variables

We then explore the distribution of the three microbiotypes among multiple clinical variables in Table 2. This shows no significant difference for some variables including asthma, aspirin sensitivity, GORD, diabetes mellitus, and current smoking status, (FDR-corrected p > 0.05; Chi-squared test). The cross tabulation however revealed a statistically significant association with “aspirin sensitivity” or aspirin-exacerbated respiratory disease (AERD) (p = 0.02), although this did not persist after a Benjamini-Hochberg correction (corrected p = 0.077). Patients who were aspirin-sensitive (or suffering from AERD) showed less prevalence of microbiotypes 1, 2 and a higher prevalence of microbiotype 3, compared to those who were not aspirin-sensitive. On the other hand, patients who were undergoing their “primary surgery”, had a higher prevalence of microbiotype 1 and a lower prevalence of microbiotype 3, compared to those patients who had had previous surgeries, but these results were not statistically significant.

**Table 2:** Distribution of microbiotypes by various clinical variables.

| variable | value | microbiotype_1 | microbiotype_2 | microbiotype_3 | p value |
| --- | --- | --- | --- | --- | --- |
| Asthma | No | 162 (56.4%) | 81 (28.2%) | 44 (15.3%) | 0.906 |
|  | Yes | 55 (51.4%) | 31 (29.0%) | 21 (19.6%) |  |
| Aspirin sensitivity | No | 202 (55.3%) | 106 (29.0%) | 57 (15.6%) | 0.077 |
|  | Yes | 12 (48.0%) | 5 (20.0%) | 8 (32.0%) |  |
| Diabetes | No | 189 (54.9%) | 98 (28.5%) | 57 (16.6%) | 0.979 |
|  | Yes | 22 (55.0%) | 11 (27.5%) | 7 (17.5%) |  |
| GORD | No | 177 (55.3%) | 93 (29.1%) | 50 (15.6%) | 0.979 |
|  | Yes | 35 (55.6%) | 17 (27.0%) | 11 (17.5%) |  |
| Current Smoker | No | 204 (54.4%) | 110 (29.3%) | 61 (16.3%) | 0.077 |
|  | Yes | 15 (57.7%) | 4 (15.4%) | 7 (26.9%) |  |
| Primary surgery | No | 92 (47.2%) | 57 (29.2%) | 46 (23.6%) | 0.114 |
|  | Yes | 130 (60.5%) | 60 (27.9%) | 25 (11.6%) |  |

### Validation of sinonasal microbiotyping on a separate dataset

We validated our approach on a separate 16S dataset we called Dataset Two. As described in the Methods section, we validated this using an independent unsupervised approach and a semi-supervised approach guided by the Main Dataset.

The first unsupervised approach yielded three clusters similar to the microbiotypes described on the Main Dataset, with one cluster exhibiting high mean relative abundance of Corynebacteria, a second cluster exhibiting high mean relative abundance of Staphylococcus, and a third cluster with other dominant genera. Plotting the first two Principal Components (Figure 4A) resulting from PCoA on the JSD matrix revealed the same triangular distribution of samples observed in Figure 1.

**Figure 4:**
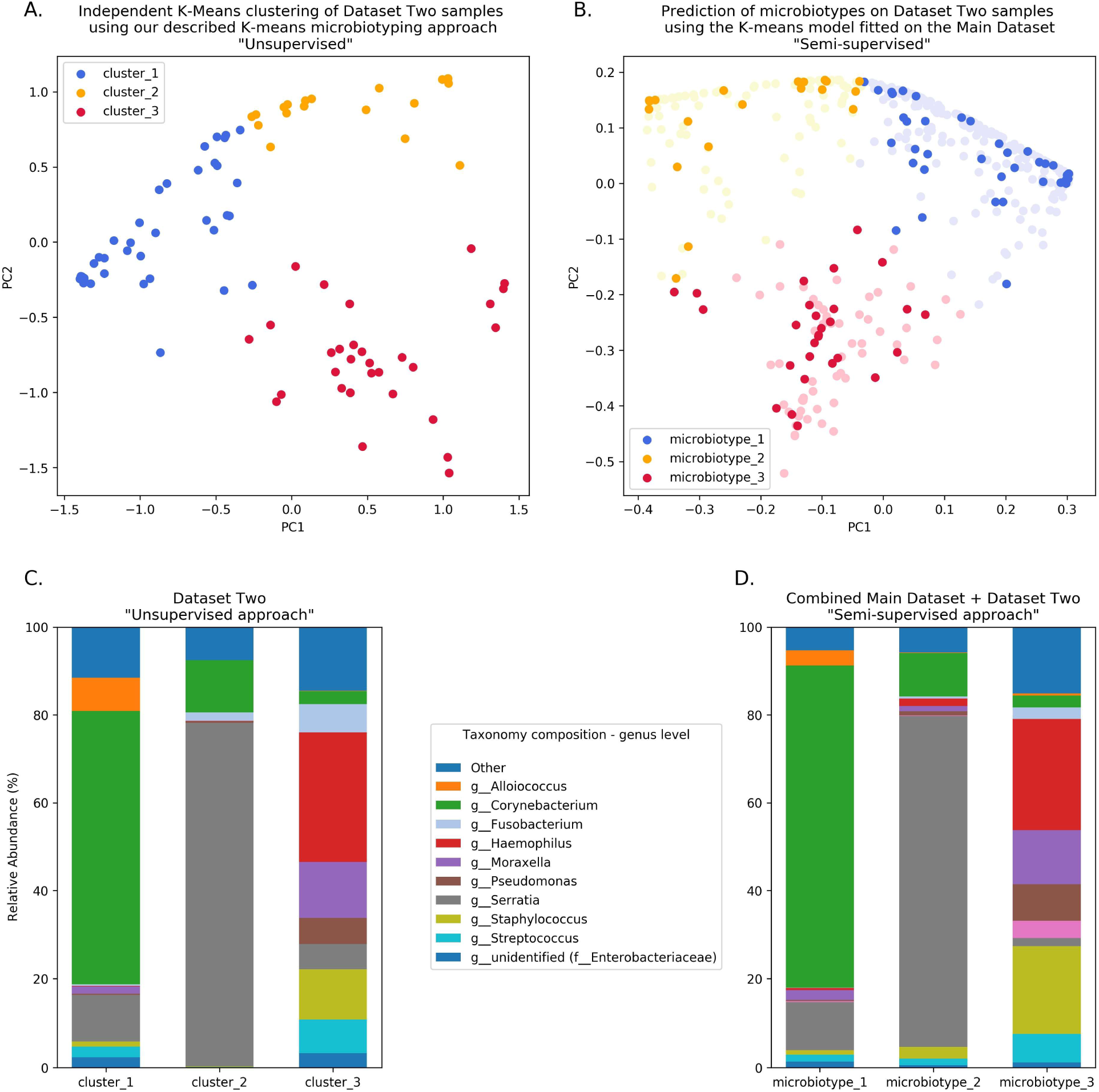
Validation of microbiotyping approach on Dataset Two.

Prevalence of the microbiotypes in this dataset (using the unsupervised approach) was as follows: microbiotype 1 assigned 39.2% of samples, microbiotype 2 with 26.8% of samples, and microbiotype 3 with 34.0%.

The second semi-supervised approach yielded similar results (Figure 4; Supplementary Table), differing in the classification of only 3 samples (out of 97 samples; i.e. 3.09%). (See Supplementary Jupyter notebook) Two of these samples show *Staphylococcus* dominating the samples in combination with *Haemophilus*, with no overt dominance of one taxon over the other, making them more-or-less transitional samples between the signatures of microbiotypes 2 and 3. The third sample was dominated by *Staphyloccocus* and *Corynebacterium*, making it a transitional sample between microbiotype 1 and microbiotype 2, with Staphylococcal species assigned to epidermidis, making this more appropriately assigned to microbiotype 1. (see Supplementary Jupyter notebook)

These results validate the microbiotyping approach and suggest that our approach and dataset could be used to guide classification of sinonasal samples sequenced in future separate studies. (Figure 4) Moreover, it points towards a potential clinical relevance of performing sinonasal microbiotyping.

## DISCUSSION

We demonstrate that the microbiota of most sinus swab samples could be classified into distinct signatures or archetypes, which we have termed “sinonasal microbiotypes”. We observed three main microbiotypes: the most prevalent being a *Corynebacterium*-dominated microbiotype (microbiotype 1), then a *Staphylococcus*-dominated microbiotype (microbiotype 2), and microbiotype 3 which includes samples dominated by *Streptococcus*, *Haemophilus*, *Moraxella*, *Pseudomonas*, and other genera (3S, 3H, 3M, 3P, and 3O respectively).

As we have previously reported,^2^ the sinus microbiota are dominated by the genera *Corynebacterium* and *Staphylococcus* (microbiotypes 1 and 2). A similar clustering approach to the sinus microbiome was applied by Cope and colleagues, who utilized Dirichlet multinomial mixture models (DMMs),^5^ and reported that most samples in their study were occupied by a continuum of Staphylococcaceae and Corynebacteriaceae.^5^ It appears that, regardless the statistical or clustering methodology utilized, it is most likely that the sinonasal microbiome consists of core organisms^2^ that have a distinct co-occurrence pattern. This could be explored through a network analysis approach and should be a future area of study.

*Staphylococcus aureus* has been perceived to be an important pathogen in sinus inflammatory disease. *Staphylococcus aureus* biofilms may act as a nidus for recurrent infections^14,15^ and as a “nemesis” of otherwise-successful sinus surgery.^16–18^ *Staphylococcus aureus* is also a producer of exotoxins, which in some cases can serve as superantigens, and these have been previously described as playing an important role in the pathogenesis of CRSwNP.^19^ *Pseudomonas aeruginosa* biofilms are also virulent organisms that are difficult to eradicate from the sinuses, and have been associated with worse clinical outcomes.^20^ Both these organisms are important pathogens in the chronic mucociliary dysfunction exhibited in cystic fibrosis. However, methicillin-resistant *Staphylococcus aureus* (MRSA) is an important nasal colonizer that could asymptomatically colonize the nose. What determines the clinical course, between asymptomatic colonization versus symptomatic pathogenicity, remains an interesting topic of research. In this study, we identified a potential reciprocal relationship between *Staphylococcus* aureus and *Corynebacterium*. Being aware of the challenges of compositional data analysis, we utilized for this purpose the specialized SparCC algorithm which infers correlations from compositional data.^21^ This finding needs to be supported by future co-culture experiments, but suggests that *Corynebacterium sp*. may be a “cornerstone” of sinus microbial health. It is important to note that our bioinformatic methodology has been intentionally designed to utilize state-of-the-art software methods at every step of the analysis pipeline, in order to maximise the resolution of taxonomy assignment.^8,9,22^ Nevertheless, our approach is still confined within the limitations of current 16S sequencing methodologies, and the confidence of assignment is reduced beyond the genus level. Our analysis pipeline could not delineate between different *Corynebacterium* at the species level and *Staphylococcus aureus* at the strain level. Hence functional difference between samples with same species remain to be determined using a functional metagenomics approach. A recent study suggest that by incorporating location information or “sample-level metadata” into species-level assignment accuracy could be improved.^23^ In our study, the differential relationships of both *Staphylococcus aureus* and *epidermidis* towards *Corynebacteria* (negative and positive associations, respectively) could be of clinical significance and is worthy of future investigation. We performed a post-hoc inspection of species-level assignment in Dataset Two, to investigate whether this finding will be reproducible in a separate dataset. This confirmed clustering of almost all *Staphylococcus aureus* species in microbiotype 2. (Supplementary Results in Jupyter Notebook)

Interestingly, we found that the distribution of the sinonasal microbiotypes was not significantly dissimilar amongst healthy controls and CRS patients. There appeared to be no significant associations with other clinical variables such as asthma and aspirin-sensitivity after controlling for multiple comparisons. (Table 2) The distribution of the microbiotypes however differed according to centre/location of collection. (Figure 3) As such, we cannot conclude based on our study that microbiotypes could function independently as a disease biomarker. Although not reaching statistical significance (chi squared p > 0.05) the prevalence of microbiotype 3 was higher in CRSsNP and CRSwNP, compared to controls. It could be the case that chronicity of inflammation -on its own-is not a determinant of a dysbiotic microbiome, but whether there is a clinically-evident “sinus infection” current at the time of sample collection. In this theory, stable chronic sinuses with no overt signs of acute or chronic infection, may remain similar to a “healthy sinus microbiome”. Only when the sinuses are clinically infected (as evident on clinical symptoms and endoscopic findings), the microbiota become disrupted and the dysbiosis exaggerated. It is important to note that *Streptococcus*, *Haemophilus* and *Moraxella* (represented here in microbiotype 3) have been traditionally implicated in acute infections of the upper respiratory tract including acute rhinosinusitis and acute otitis media. Unfortunately, information regarding acute exacerbations was not explored within this study.

Regarding geographical differences: Asia and Australasia showed an over-representation of microbiotype 1. Europe had a higher prevalence of microbiotype 2. Unfortunately, the study only included one European centre (Amsterdam) so it is difficult to be certain whether this finding generalizes to other locations in Europe. The driving factors for these geographical differences could be multiple, including but not limited to clinical practices such as local antibiotic prescriptions for CRS and timing of recruitment of patients for sinus surgery, as discussed previously.^2^

We have adapted our methodology from the enterotyping approach taken by Arumugam et al.^4^ for classifying bacterial signatures of the gut microbiome. In their original manuscript, they described three different enterotypes in the gut dominated by *Prevotella*, *Bacteroidetes*, and *Ruminococcus* respectively.^4^ Several papers have correlated gut enterotypes with various clinical variables.^24,25^ Despite this, enterotyping as an approach to population stratification has not been without its controversies. Several authors have criticized the definition of distinct clusters, since it neglects intra-cluster variation and gradients between clusters.^26–29^ We provide answers to previous critique^28^ to enterotyping as it applies to our study in Supplementary Table S2. It is important to note these valid criticisms to any community typing approach. In our experiment, the clusters or types lie on a continuum, with some samples falling in the gradients between two, or perhaps even all three microbiotypes (see ordination plots). The histograms in Figure 2 also suggest this, but they do show most samples in each microbiotype feature a high relative abundance of a dominating genus in many samples. We investigated a simple dominance measure, the Berger-Parker (BP) alpha diversity index,^30^ in the combined datasets’ 507 samples. The Berger-Parker index simply reports the relative abundance of the most dominant taxon in a sample. This found that only 24.9% of samples had a dominating taxon that only had a relative abundance of 50% or less. On the other hand, 51.9% of samples had the dominant taxon exhibiting a relative abundance of greater than 70% of the sample.(Supplementary Results in Jupyter notebook; Supplementary Figure S1) This shows that in most samples, there is one dominating organism. Based on these results, the microbiotyping approach is therefore proposed to reduce complexity about modeling bacterial interactions in the sinuses, and not to suggest that each type is a walled-off discrete cluster. Further investigations into the local substructures of each type will be required to further explore the roles and interactions of its constituent taxa. Another limitation of our description of microbiotypes is that they may as well describe different community “states” rather than community “types”, since we do not have longitudinal data to describe how these clusters behave with the passage of time and treatments. Hence, we could not confirm whether these are stable, consistent communities across time. It may well be that intermediate samples lying between two or more microbiotypes are representing a transitional state. An important future avenue of research is to conduct a longitudinal study to investigate the temporal stability of these clusters.

We predict that ongoing sinonasal microbiome research and consequent large meta-analyses of microbiota studies, with novels tools (such as QIITA^31^) enabling such large-scale studies, will allow the refinement of these types and further clarify their clinical/microbiological utility. Our methodological approach to describe the microbiotypes is not exclusive, as alternative statistical or machine-learning approaches could be employed to investigate them. In light of this, we expect that international multi-centre standardization and rationalization of the sinonasal microbiotypes would be possible in the future, similar to the recent proposed effort to standardize enterotyping of the gut microbiota by Costea et al.^29^

## CONCLUSION

We investigated the ISMS dataset through an approach modeled on human gut microbiome enterotyping and we found three major microbial community types or “microbiotypes” as clusters that lie on a continuum, based on an unsupervised machine learning approach that involved dimensionality reduction and clustering. Microbiotypes did not show an association with disease state or clinical variable, suggesting that they could not function as independent disease biomarkers. The description of these microbiotypes has also unveiled a potential reciprocal relationship between *Staphyloccocus aureus* and *Corynebacterium spp*. in the sinuses that requires further investigation in future studies. The findings were validated on a separate previously unpublished sinus bacterial 16S gene dataset. Microbiotypes are therefore proposed to reduce the complexity of modeling bacterial interactions in the sinuses, and in this sense hold microbiological and clinical relevance that could potentially influence medical and surgical treatment of CRS patients.

## METHODS

### The “International Sinonasal Microbiome Study (ISMS)” dataset

We perform the primary analysis on the dataset obtained from the “International Sinonasal Microbiome Study (ISMS)” project.^2^ In summary, this dataset is a multi-centre 16S-amplicon dataset which includes endoscopically-guided, guarded swabs collected from the sinuses (in particular the middle meatus / anterior ethmoid region) of 532 participants in 13 centres representing 5 continents. Details of sample collection, DNA extraction and sequencing methodologies are described in the original report.^2^ The 16S gene region sequenced was the V3–V4 hypervariable region, utilizing primers (CCTAYGGGRBGCASCAG forward primer) and (GGACTACNNGGGTATCTAAT reverse primer) according to protocols at the sequencing facility (the Australian Genome Research Facility; AGRF). Sequencing was done on the Illumina MiSeq platform (Illumina Inc., San Diego, CA) with 300-base-pairs paired-end Illumina chemistry

### Bioinformatics pipeline

Details of the bioinformatic pipeline is detailed in the original report.^2^ In summary, we utilized a QIIME 2-based pipeline.^8^ Forward and reverse fastq reads were joined^32^, quality-filtered,^33^, abundance-filtered^34^, then denoised using deblur^9^ through QIIME 2-based plugins. This yielded a final feature table of high-quality, high-resolution Amplicon Sequence Variants (ASVs). Taxonomy assignment and phylogenetic tree generation^35^ was done against the Greengenes^36^ database; and taxonomy was assigned using the QIIME 2 BLAST assigner.^22^ A rarefaction minimum depth cut-off was chosen at 400 and this yielded 410 samples out of the original 532 for downstream analysis. The same pipeline was then applied on DataSet Two for purposes of validation of microbiotyping. We chose to reproduce exactly all the original pipeline steps on DataSet Two, despite being a completely separate dataset, to reduce bias.

### Delineating the microbiotypes of the sinonasal microbiome

Our approach was guided by the “enterotyping” method described by Arumugam et al.^4^ with adaptations. We constructed a sample distance matrix using the Jensen-Shannon distance (JSD) metric, as used in the original “enterotypes” paper.^4^ The Jensen-Shannon distances were calculated between samples in the genus-level-assigned table in a pairwise fashion using the JSD function in the R package “philentropy” with a log (log_10_) base. Following this, Principal Coordinate analysis (PCoA) was done on the distance matrix for dimensionality reduction and visualization. Clustering was then performed using a standard K-means clustering algorithm, as implemented in the machine learning Python package scikit-learn (version 0.20.1);^37^) on the first two principal components (PCs) obtained from the PCoA, with the number of clusters (k) chosen at 3 based on visual inspection of the beta diversity PCoA plots. Average silhouette scores, as implemented in scikit-learn, for the range (k = 2 - 8) were calculated to assess clustering quality, and this revealed the highest silhouette scores: 0.61 and 0.6 for [k=4] and [k=3] respectively. The three resulting clusters were defined as the three sinonasal microbiotypes. For further exploration of the subgroups that constitute microbiotype 3, we used the hierarchical density-based clustering algorithm “hdbscan”^7^ on the full-dimensional feature table. Genera were projected onto the PCoA matrix using a biplot approach^6^, as implemented in scikit-bio’s function *“pcoa_biplot”*. Genera were represented in the biplot figure as arrows, originating from the centre of the plot pointing to the direction of the projected feature coordinates, and the lengths normalized as a percentage of the longest arrow. We utilized “Analysis of Compositions of Microbiomes (ANCOM)”^38^ for identifying differentially-abundant taxa. Taxa genus level and Staphylococcus species level co-occurrence/correlation analysis were done after taxonomy assignment using SparCC algorithm,^21^ in the fast implementation in FastSpar.^39^

### Validating microbiotypes on a second sinonasal microbiome dataset

To infer whether our classification could be generalizable to other sinonasal microbiome samples not included in this study, we sought to validate our microbiotyping approach on a separate, previously-unpublished, 16S dataset. This dataset includes sinonasal microbiome swabs collected from private and public patients attending the Otolaryngology Department (University of Adelaide) to have surgery done by the authors P.J.W., A.J.P. or the Otorhinolaryngology Service at the Queen Elizabeth Hospital in Adelaide, South Australia. Similar to the main dataset, these included CRS patients who underwent endoscopic sinus surgery for this sinus disease, and non-CRS control patients who underwent other otolaryngological procedures, such as tonsillectomy, septoplasty or skullbase tumour resection. Sample collection, and processing were done in a standardized fashion similar to that has been described in the ISMS main dataset, except that DNA extraction was carried out using the PowerLyzer Power-Soil DNA kit (MoBio Laboratories, Salona Beach, CA) as previously described^40^, rather than the Qiagen DNeasy kit (Qiagen, Hilden, Germany). Similar to the ISMS samples, library preparation and 16S sequencing were done at the Australian Genome Research Facility (AGRF) on the Illumina MiSeq platform (Illumina Inc., San Diego, CA, USA) with the 300-base-pairs paired-end chemistry. Libraries were generated by amplifying (341F–806R) primers against the V3–V4 hypervariable region of the 16S gene (CCTAYGGGRBGCASCAG forward primer; GGACTACNNGGGTATCTAAT reverse primer).^41^ PCR was done using AmpliTaq Gold 360 master mix (Life Technologies, Mulgrave, Australia) following a two-stage PCR protocol (29 cycles for the first stage; and 8 cycles for the second, indexing stage). Sequencing was done over two MiSeq runs in January 2015. We termed this dataset in this manuscript “Dataset Two”. This dataset comprises samples collected from 129 participants. Rarefaction at a cutoff of 400 reads was performed, to match what was performed for the main dataset, and samples with read number less than 400 were excluded; this yielded a final feature table containing 97 samples, representing 33 CRSsNP patients, 35 CRSwNP patients, and 29 controls.

We took two separate approaches to validation. The first approach is to replicate the previously-described unsupervised K-means microbiotyping methodology independently on samples in Dataset Two. We call this first approach the “unsupervised approach”. The second approach is to use the K-means model that was fitted on the samples from the Main Dataset to predict labels (i.e. microbiotypes) of the samples in Dataset Two. As such, the Main Dataset is used as a “training dataset” in the language of machine learning. We called the second approach the “semi-supervised approach”.

### Statistical Analysis

All frontend analyses were done using the Jupyter notebook frontend^42^ and utilizing the assistance of packages from the Scientific Python^43^ stack (numpy, scipy, pandas, statsmodels), scikit-learn^37^, scikit-bio (https://github.com/biocore/scikit-bio) and omicexperiment (https://www.github.com/bassio/omicexperiment).

---

**Figure S1:**
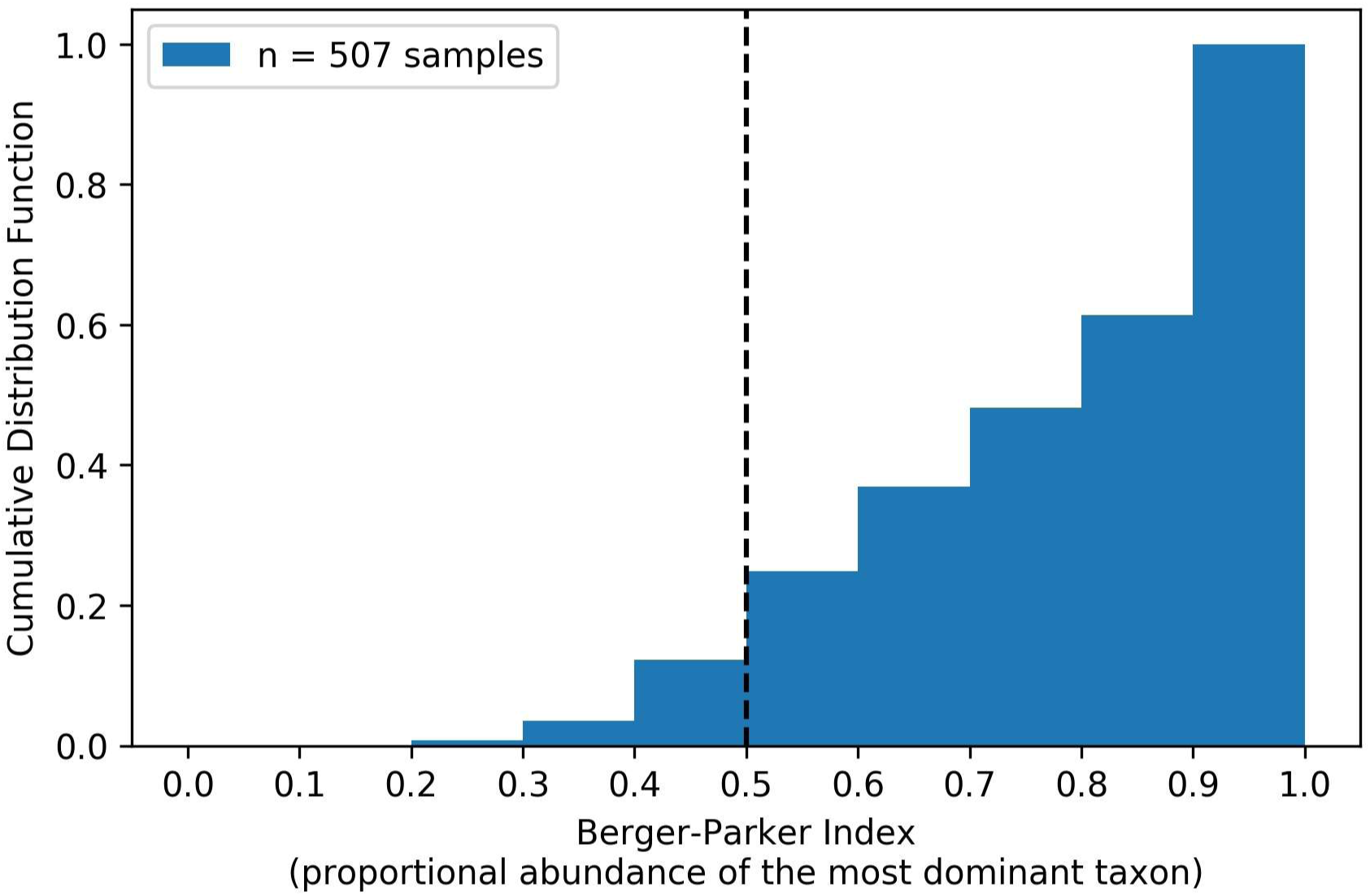
Cumulative distribution function of the Berger-Parker Index in the combined datasets.

**Table S1A:** Predominant taxa of microbiotype 1.

| genus | Mean Relative Abundance (%) | Prevalence (%) |
| --- | --- | --- |
| Corynebacterium | 75.29 | 100 |
| Staphylococcus | 10.69 | 76.58 |
| Alloiococcus | 2.79 | 28.83 |
| Moraxella | 2.31 | 9.91 |
| unidentified<br>(Enterobacteriaceae) | 1.41 | 15.32 |
| unidentified (Neisseriaceae) | 1.18 | 20.72 |
| Streptococcus | 1 | 21.62 |
| Haemophilus | 0.56 | 9.91 |
| unidentified (Moraxellaceae) | 0.44 | 2.7 |
| Ralstonia | 0.34 | 10.36 |

**Table S1B:**
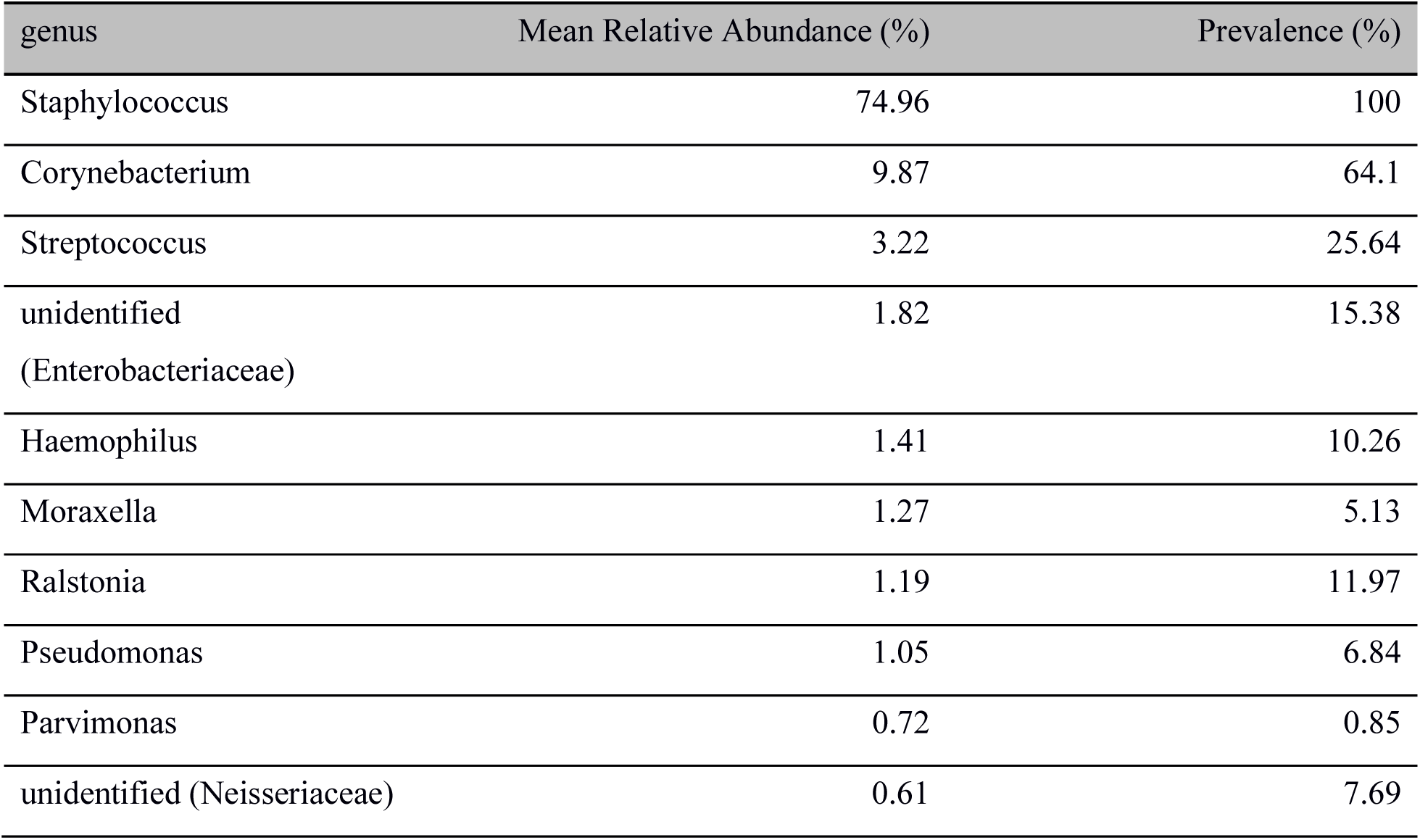
Predominant taxa of microbiotype 2.

**Table S1C:**
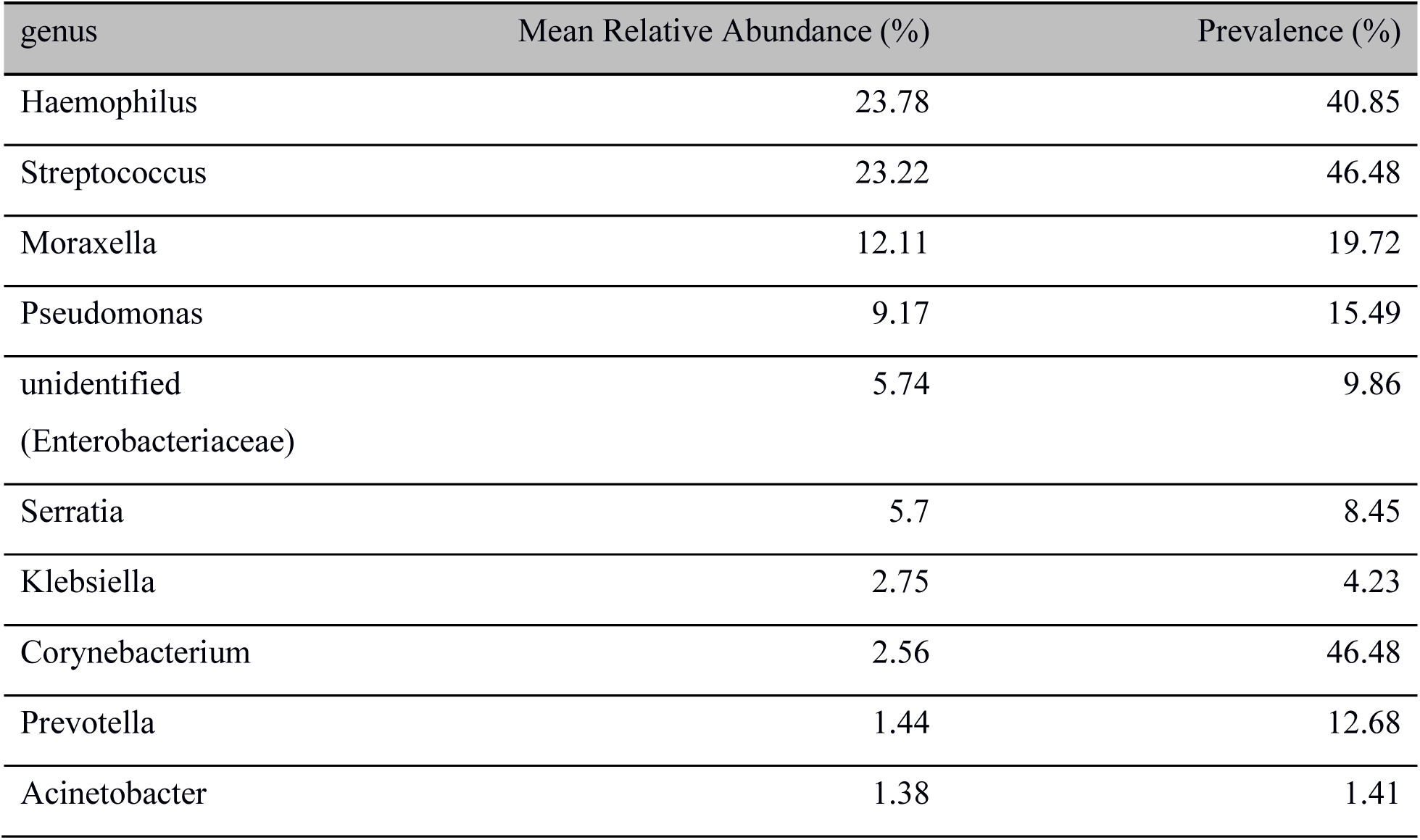
Predominant taxa of microbiotype 3.

**Table S2:** Addressing previous criticism to gut enterotyping.

| Critique | Answer |
| --- | --- |
| Discrete clusters or a multi-dimensional gradient? | We acknowledge the a proportion of samples fall in the gradient between the proposed microbiotypes. Berger-Parker index investigation showed that most samples had one dominating taxon. |
| Do discrete clusters link to human disease? | No. We report that we could not find an association between the microbiotype and chronic sinusitis disease status. |
| Is sampling frame or selection bias affecting results? | No; Multi-centre international study with consecutive sampling methodology. We also validate on a separate dataset. |
| Use inappropriate visualization such as “star-burst plots”? | We did not use inappropriate visualizations. |
| Use a supervised approach “between-class analysis”? | We use an unsupervised clustering and dimensionality reduction approach. |
| Is an individual’s microbiotype stable over time? | Answer unknown; Future longitudinal studies required. |

